# Molecular Mechanism of Condensin I Activation by KIF4A

**DOI:** 10.1101/2024.03.19.585759

**Authors:** Erin E. Cutts, Damla Tetiker, Eugene Kim, Luis Aragon

## Abstract

During mitosis, the condensin I and II complexes compact chromatin into chromosomes. Loss of the chromokinesin, KIF4A, results in reduced condensin I chromosome association. However, the molecular mechanism behind this phenotype is unknown. Here, we show that KIF4A binds directly to condensin I HAWK subunit, NCAPG, via a conserved disordered short linear motif (SLiM) found in its C-terminal tail. KIF4A binding directly competes with two auto-inhibitory interactions in condensin I, mediated by SLiMs found in the N-terminus of NCAPH and the C-terminus of NCAPD2, which bind sites that overlap with KIF4A binding on NCAPG. Addition of the KIF4A SLiM peptide alone is sufficient to stimulate condensin’s ATPase and DNA loop extrusion activity. We also identify SLiMs in known yeast condensin interactors, Sgo and Lrs4, that bind yeast Ycg1, the HAWK equivalent to NCAPG. Our findings, together with previous work on condensin II and cohesin, demonstrate that SLiMs binding to HAWK subunits is a conserved mechanism of regulation in SMC complexes.

## Introduction

Ensuring faithful division of genetic material between daughter cells is a challenge every cell faces during mitosis. Cells must solve two important problems before attempting genome separation; (i) interphase chromosomes are generally much longer than the dividing cell, hence need to be compacted, and (ii) replication causes entanglement between the newly produced sister chromatids, hence these need to be resolved ahead of anaphase^1^. To overcome these two issues, cells organise interphase chromatin into compact rod-shaped chromosomes during mitosis, reducing their length sufficiently so that chromosome entrapment during cytokinesis is avoided and promoting the disentanglement of sister chromatids arms.

In mammalian cells, the key players driving the transformation of interphase chromatin into mitotic chromosomes are two structural maintenance of chromosome (SMC) family members; condensin and cohesin. Condensin creates DNA loops^2, 3^ around an axis to facilitate chromosome compaction^4^, while cohesin provides chromosome “cohesion” by holding chromatids together^4–6^. Cohesin also has DNA looping activity^7, 8^ restricted to interphase^4^ which enables long-range genome interactions^9^, in contrast to condensin, which only acts during mitosis.

Condensin and cohesin create loops by loop extrusion, where DNA loop size is processively increased in an ATP-dependent manner. This mechanism is conserved through evolution, with all SMC complexes shown to have loop extrusion activity *in vitro*^2, 3, 7, 8, 10, 11^, however a full molecular understanding as to how loop extrusion is achieved and regulated remains elusive. This conserved activity in the condensin and cohesin SMC complexes is likely due to a conserved overall architecture, composed of heterodimers of SMC subunits, a kleisin subunit, and two different HAWK (HEAT associated with Kleisin) proteins (Figure 1A). SMC subunits have ∼50 nm long coiled-coil regions, which heterodimerise via a hinge domain at one end of the coiled coils, and sandwich two ATP molecules between split ATPase domains at the other end. The kleisin binds all components of the complex; its N-terminus associates with one SMC, its C-terminus with the other SMC, and the middle section binds both HAWK proteins^12^. Current models for the loop extrusion mechanism suggest each HAWK subunit helps to define compartments that can hold and exchange DNA^10, 13–15^. For condensin, the HAWK bound towards the C-terminus of the Kleisin, Ycg1/NCAPG is thought to anchor the complex to DNA, with structural work demonstrating that the Kleisin, Brn1/NCAPH, folds and latches over DNA^16, 17^. While the other HAWK, Ycs4/NCAPD2, along with the Kleisin and ATPase heads, defines another compartment necessary for DNA loop growth^15, 18^.

**Figure 1:**
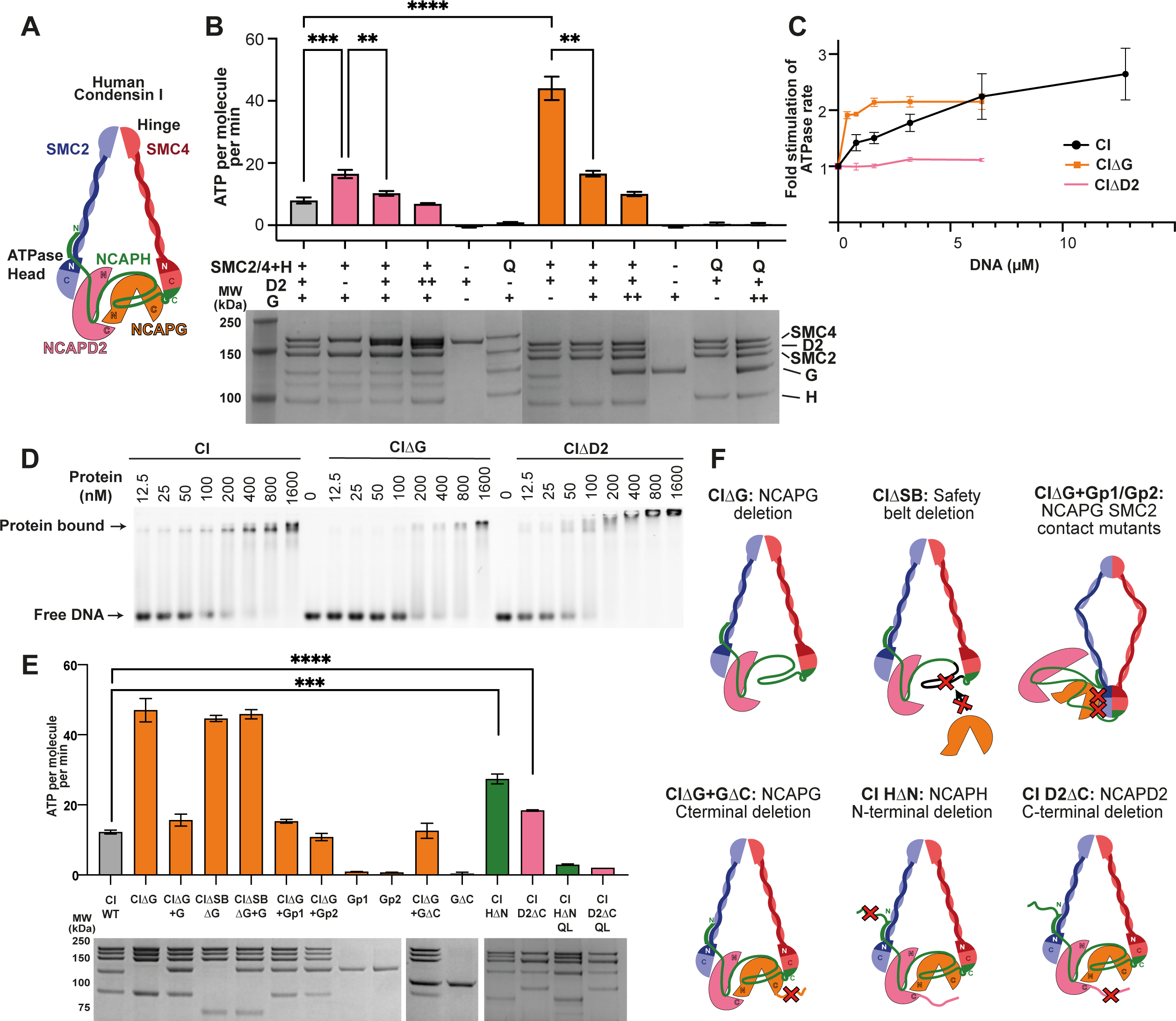
Condensin I mutations results in increased ATPase activity. (A) An overview of the subunit architecture of human condensin I. (B) ATPase assay of human condensin I and II, respectively, lacking HAWK domains. + indicates subunit is present is a stoichiometric amount, while ++ indicates it is present in a 2-fold excess. Q refers to complexes where there is a mutation in the Q-loop which are unable to bind ATP. The lower panel are examples of SDS page gels of the ATPase reaction mixture after completion of the reaction. (C) Effect of 50 bp dsDNA on ATPase rate of condensin I. (D) Gel shift assay of condensin I and II pentamers and tetramers, respectively, binding to 50 bp dsDNA labelled with cy5. (E) ATPase assay of condensin I mutations shown in (F). Mutation CIΔG indicate tetrameric condensin I lacking NCAPG. CIΔSB indicates deletion of the NCAPH region where NCAPG binds. Gp1 and Gp2 indicates mutations in NCAPG homologous to patch 1 and 2 where Ycg1 and SMC2 interact in yeast condensin, GΔC indicates deletion of NCAPG C-terminal disordered region, CINΔH indicates deletion of NCAPH N-terminal disordered region and CID2ΔC indicates deletion of NCAPD2 C-terminal disordered region, CIQHΔN and CIQD2ΔC indicating ATPase dead Q-loop mutant controls. All ATPase assays from at least 3 repeats, error bars indicate standard error, two tail t-test *p<0.05, **p<0.01, ***p<0.001, **** p<0.0001.

How condensin loop extrusion activity is controlled by the cell cycle is a long-standing question. While previous studies have demonstrated phosphorylation by mitotic kinases such as CDK1/cyclin B and Aurora contribute to activation^19^, mechanistic detail is lacking. Curiously, recent studies have demonstrated that chromosomes can still be produced by tetrameric condensin I or II lacking the NCAPG or G2 subunit, respectively^20, 21^, suggesting that while tetrameric condensins (lacking NCAPG/G2 subunits) are capable of compaction, they lack fine-tuned regulation of their activity. HAWK subunits NCAPG2 and NCAPG have binding sites for other proteins that regulate the activity of condensins. NCAPG2 acts as a binding scaffold for a Short Linear Motif (SLiM) found in MCPH1, whose binding results in condensin II repression^22, 23^. While the chromokinesin, KIF4A, is known to bind NCAPG and is a regulator that promotes the association of condensin I to chromosomes as well as the congregation of chromosomes in metaphase^24–27^. However, the molecular mechanisms of how KIF4A regulates condensin I activity has not been determined.

In this work, we investigate the contribution of condensin I HAWK subunits to the core ATPase activity of the complex. We demonstrate that NCAPG acts as a scaffold for three SLiMs; two resulting in auto-inhibition, found in C- and N-terminal disordered regions of NCAPD2 and NCAPH, respectively, and one in KIF4A resulting in activation. We show that competition between KIF4a and NCAPD2/H interactions results in condensin I activation. Importantly, we find SLiM mediated regulation is a feature that NCAPG shares with other HAWK subunits like Ycg1 from yeast condensins, NCAPG2 from condensin II, and STAG2 from cohesin.

## Results

### Human Condensin Tetramers have Hyperactive ATPase Activity

To test the effect HAWK subunits have on the core activity of condensin, human wild-type condensin I pentameric complexes (CI) as well as tetrameric complexes lacking either NCAPG (CIΔG) or NCAPD2 (CIΔD2) were purified and assayed for ATPase activity (Figure 1B). CIΔG and CIΔD2 hydrolysed ATP significantly faster, ∼4 times and ∼2 times, respectively, than CI. The hyperactivity of tetrameric complexes was rescued to near pentameric levels when an excess of recombinant NCAPG or NCAPD2 were added, with the addition reconstituting the pentameric complex (Supplementary Figure 1A and B). CIΔG and CIΔD2 with mutations in the Q-loop of ATPase site, unable to bind ATP, (CIΔG-Q and CIΔD2-Q) and NCAPG or NCAPD2 had negligible ATPase activity, suggesting the ATPase activity measured is indeed that from the condensin I active site. Adding excess NCAPG or NCAPD2 to pentameric CI had no effect, demonstrating that super stoichiometric HAWK subunits did not further enhance repression (Figure 1B). Hence, NCAPG and NCAPD2 both have roles in repressing condensin I complex ATPase activity.

### NCAPD2 is required for DNA stimulated ATPase, while NCAPG represses it

The ATPase activity of condensin complexes is known to be stimulated by DNA and structural data demonstrates both HAWK domains bind DNA^15, 16, 18^. To determine how each HAWK domain affects DNA stimulated ATPase rates we measured ATPase activity while titrating in 50 bp dsDNA. Under these conditions, CI ATPase rate is stimulated up to a limit of ∼2.4-fold at saturating DNA concentrations (Figure 1C). DNA stimulation was completely abolished in tetrameric condensin lacking NCAPD2 CIΔD2 (Figure 1C). This is consistent with previous studies using yeast condensin which demonstrate the NCAPD2 equivalent subunit, Ycs4, clamps DNA on top of the ATPase heads to stimulate activity^15^. In contrast, tetrameric complexes lacking NCAPG, CIΔG, were maximally stimulated at ∼10-fold lower DNA concentrations than pentameric CI, suggesting that NCAPG also represses DNA stimulated ATPase activity (Figure 1C).

To rule out that differences in DNA binding affinity between pentameric CI and tetrameric CIΔG contribute to the difference in the observed ATPase stimulation, we examined the binding of tetrameric and pentameric complexes to DNA using gel shift assays (Figure 1C). While loss of NCAPD2 resulted in a minimal reduction in binding, loss of NCAPG resulted in ∼4-fold reduction in affinity (Figure 1D); therefore CIΔG’s hypersensitivity to DNA is not due to an increase in DNA binding affinity but rather due to NCAPG mediated repression of ATPase activity in response to DNA.

### Mechanisms of NCAPG-derived repression

Our data demonstrates that the presence of HAWKs in condensin complexes represses their core ATPase activity (Figure 1B and C). We hypothesised three potential mechanisms: 1) NCAPG binds to the kleisin NCAPH via an extensive interface composed of ∼120 amino acids of NCAPH (residues ∼400-520). Hence loss of NCAPG could potentially alleviate structural constraints by lengthening the kleisin, resulting in higher ATP turnover. 2) A direct interaction between the yeast NCAPG homologue, Ycg1, and SMC2 was observed in the structure of ATP bound condensin^28^, raising the possibility that this might inhibit complex activity. 3) Recent work has suggested that disordered regions in non-SMC subunits of condensin I and II can act as regulators of activity during chromosome assembly^21, 29^. Therefore it might be that the absence of NCAPG alters the regulatory roles of these regions, causing higher ATPase rates.

To further investigate these hypotheses, we generated a panel of mutations and deletions (Figure 1E and F). First, we generated an internal deletion in NCAPH extending from residues 421-539 (CIΔSB) that covers the region of interaction with NCAPG. This deletion did not rescue the increased ATPase rate observed for tetrameric CIΔG, suggesting that the potential structural constraint on the kleisin imposed upon NCAPG binding is not behind the increased ATPase rate of CIΔG. Next, we tested whether mutating the predicted interface between NCAPG and SMC2 would recapitulate the increase in ATPase rate observed. Based on previous Ycg1/SMC2 interface mutations^28^ we generated two groups of mutations, NCAPGpatch1 and NCAPGpatch2. The ATPase rates for both mutant proteins exhibited comparable rates to wild-type pentameric CI, suggesting that NCAPG/SMC2 interactions do not suppress activity. Finally, we tested whether deletion of disordered regions at either the C-terminus of NCAPG (CIGΔC) or NCAPD2 (CID2ΔC), or the N-terminus of NCAPH (CIHΔN) resulted in elevated ATPase activity. While CIGΔC was not significantly different from wild-type, CID2ΔC exhibited a 1.5-fold increase in ATPase activity while CIHΔN resulted in a 2.3-fold increase (Figure 1E). These results show that disordered regions in NCAPD2 and NCAPH reduce the core ATPase rate of condensin I.

### Direct interaction between SLiMs in NCAPD2 and NCAPH with NCAPG regulates complex activity

Next, we sought to explore the mechanisms by which the disordered regions of NCAPD2 and NCAPH might affect the ATPase activity. Previous studies have shown that a weak interaction may exist between NCAPD2/D3 and G/G2^3, 12, 30^. We therefore considered the possibility that interactions involving NCAPG and the disordered regions identified on NCAPD2 and NCAPH could be involved in the regulation of ATPase activity, particularly since CIΔG exhibited the strongest increase in ATPase rate (Figure 1B).

We ran AlphaFold2 multimer predictions of these regions with NCAPG ^31, 32^, resulting in predictions with high-confidence pLDDT scores and low predicted alignment error in both cases (Figure 2A and D, Supplementary Figure 2A and B). In both cases the interface is mediated by Short Linear Motifs (SLiMs) in the disordered regions binding a structured scaffold on NCAPG. The interface between the NCAPD2 SLiM and NCAPG is composed of conserved basic (residues 1393-1400) and acidic (residues 1372-1383) patches, and I1391 which is buried in a hydrophobic pocket on NCAPG (Figure 2A and B, Supplementary Figure 2A). The interface between NCAPH SLiM and NCAPG occurs at a different region of NCAPG via conserved acidic and basic patches spanning residues 59-72 of NCAPH (Figure 2D and E, Supplementary Figure 2B).

**Figure 2:**
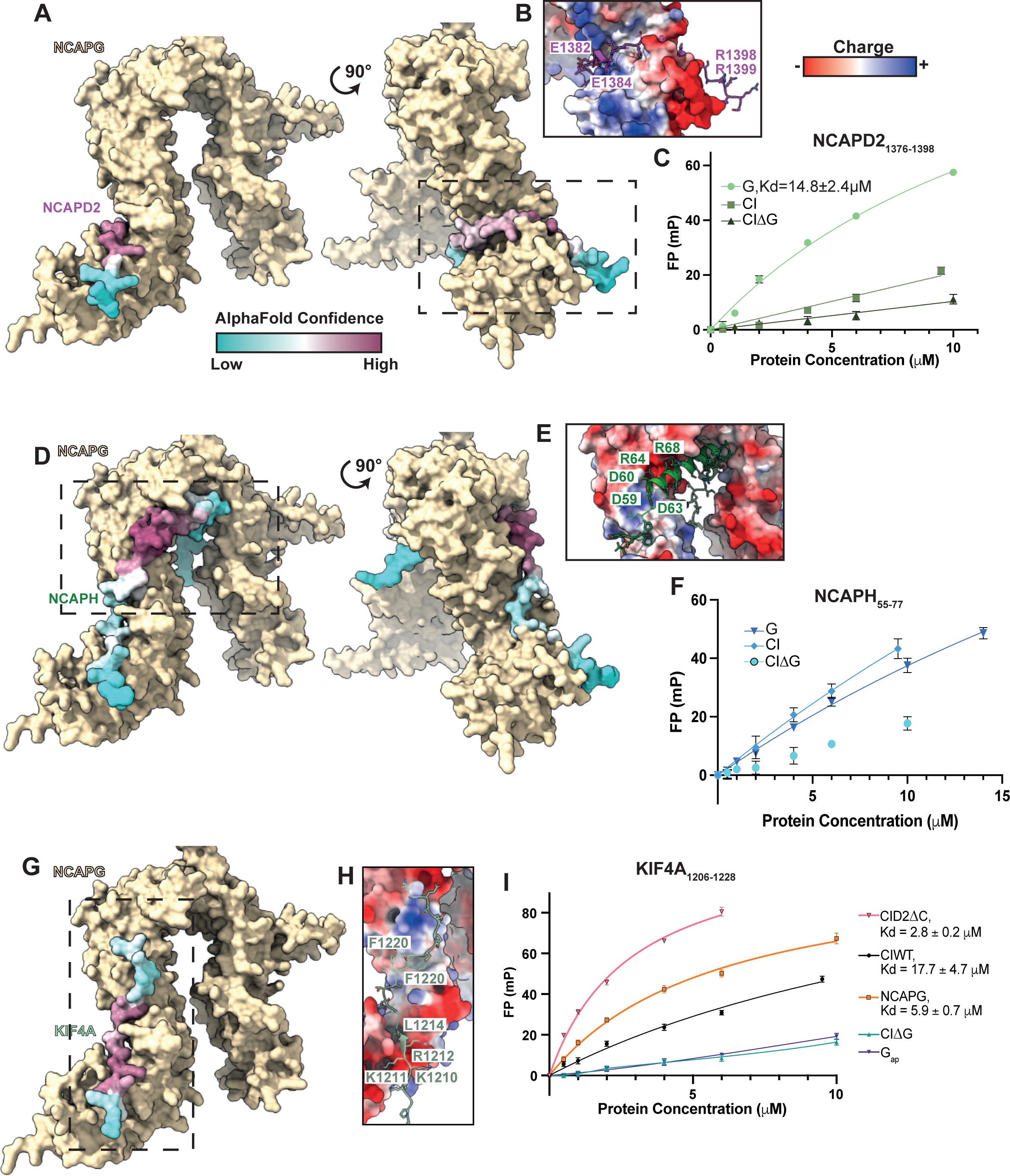
NCAPG binds disordered SLiMs regions of NCAPD2, NCAPH and KIF4A. (A) AlphaFold2 model of NCAPG with the C-terminal end of NCAPD2, with NCAPD2 coloured with AlphaFold2^31, 32^ pLDDT confidence score. (B) Surface charge of the NCAPG interface. (C) Fluorescence polarisation binding assay of 5-FAM labelled NCAPD2. (D), (E) and (F) Equivalent data for NCAPH. (G), (H) and (I) Equivalent data for KIF4A, Gap indicates NCAPG with acid patch mutation (D136K, D137K, D141K). FP data all from at least three repeats, error bars indicate standard error. Fit K_d_ shown ± standard error.

To validate the interactions predicted by AlphaFold2, we tested whether peptides containing these regions would interact with purified NCAPG and condensin I. To this aim, we used fluorescent polarisation (FP) binding assays using 5FAM labelled peptides composed of residues 1376-1398 of NCAPD2 or 55-77 of NCAPH. Titrating in increasing concentration of NCAPG resulted in a fit Kd of 14.8 ± 2.4 μM for the NCAPD2 peptide (Figure 2C). Titrating in condensin I lacking the NCAPG subunit resulted in no detectable binding (Figure 2C), consistent with specific binding of the peptide to NCAPG. Titrating in condensin I also resulted in less binding than NCAPG alone, suggesting the NCAPD2 peptide was not able to compete with the same region tethered to the complex (Figure 2C). The NCAPH peptide resulted in an increase in FP signal when titrating either NCAPG alone or Condensin I pentamers, as compared to Condensin tetramer lacking NCAPG, suggesting binding, albeit with low affinity (Figure 2F). Collectively, these results support the interaction predicted by AlphaFold2.

### Condensin I inhibitory Peptides Compete with KIF4A

Our results indicate that human condensin I is partially auto-inhibited, hence it could be activated *in vivo* by overcoming these interactions. In cells, condensin I is known to interact directly with the chromosome-associated kinesin, KIF4A, which is necessary for condensin I chromosome association^24, 25, 33^. Next, we sought to investigate the potential location of KIF4A binding on NCAPG in relation to the auto-inhibitory interaction. AlphaFold2 analysis resulted in a high-confidence prediction for an NCAPG interaction with a conserved SLiM in the C-terminal tail of KIF4A (Figure 2G and H). Strikingly, the predicted KIF4A binding sites partially overlapped with that bound by NCAPD2 and NCAPH, (Figure 2A and D). The KIF4A interaction was mediated by a conserved basic region spanning residues 1208-1212, a buried hydrophobic, L1214, which bound to the same site on NCAPG as NCAPD2, a hydrophobic region including F1220 and 1221 and a lower confidence acidic region which binds the same region as the acidic patch of NCAPH. The basic region and F1220/1221 have been demonstrated to be required for KIF4A to bind condensin II in cells^25, 33^.

To validate the AlphaFold2 model we used fluorescence polarisation assays with a 5-FAM labelled KIF4a peptide spanning residues 1206-1228. This KIF4A peptide bound to NCAPG with a Kd of 5.9 ± 0.7 μM, while no binding was detected to CIΔG, which lacks the KIF4A binding site, nor to NCAPG harbouring mutations in the acidic patch which the KIF4A basic region is predicted to bind (Figure 2I). KIF4A had a weaker affinity of 17.7 ± 4.7 μM for pentameric condensin I containing the NCAPD2 and NCAPH SLiMs, but bound with a Kd of 2.8 ± 0.2 μM to condensin I pentamer, lacking the C-terminal region of NCAPD2 (CID2ΔC), suggesting KIF4A competes with NCAPD2 to bind NCAPG.

### KIF4A activates Condensin I ATPase activity

Given that KIF4A competes with auto-inhibitory interactions, we hypothesised that KIF4A might activate condensin I ATPase activity. We assayed the ATPase activity of pentameric condensin I complex in the presence and absence of the KIF4A peptide and found a small but significant increase in ATPase activity (Figure 3A). As KIF4A bound CID2ΔC with higher affinity, we repeated the ATPase assay using CID2ΔC and found a larger effect (Figure 3A). Importantly addition of a KIF4A peptide harbouring mutations in the predicted interface, K1210A, K1211A, R1212A, L1214G, S1216A, N1217A (KIF4A mut) resulted in no increase in activity (Figure 3A). This data is consistent with KIF4A activating condensin I, and importantly, this effect is observed in the absence of NCAPD2 C-terminus, suggesting concerted action of NCAPH, NCAPD2 and KIF4A.

**Figure 3:**
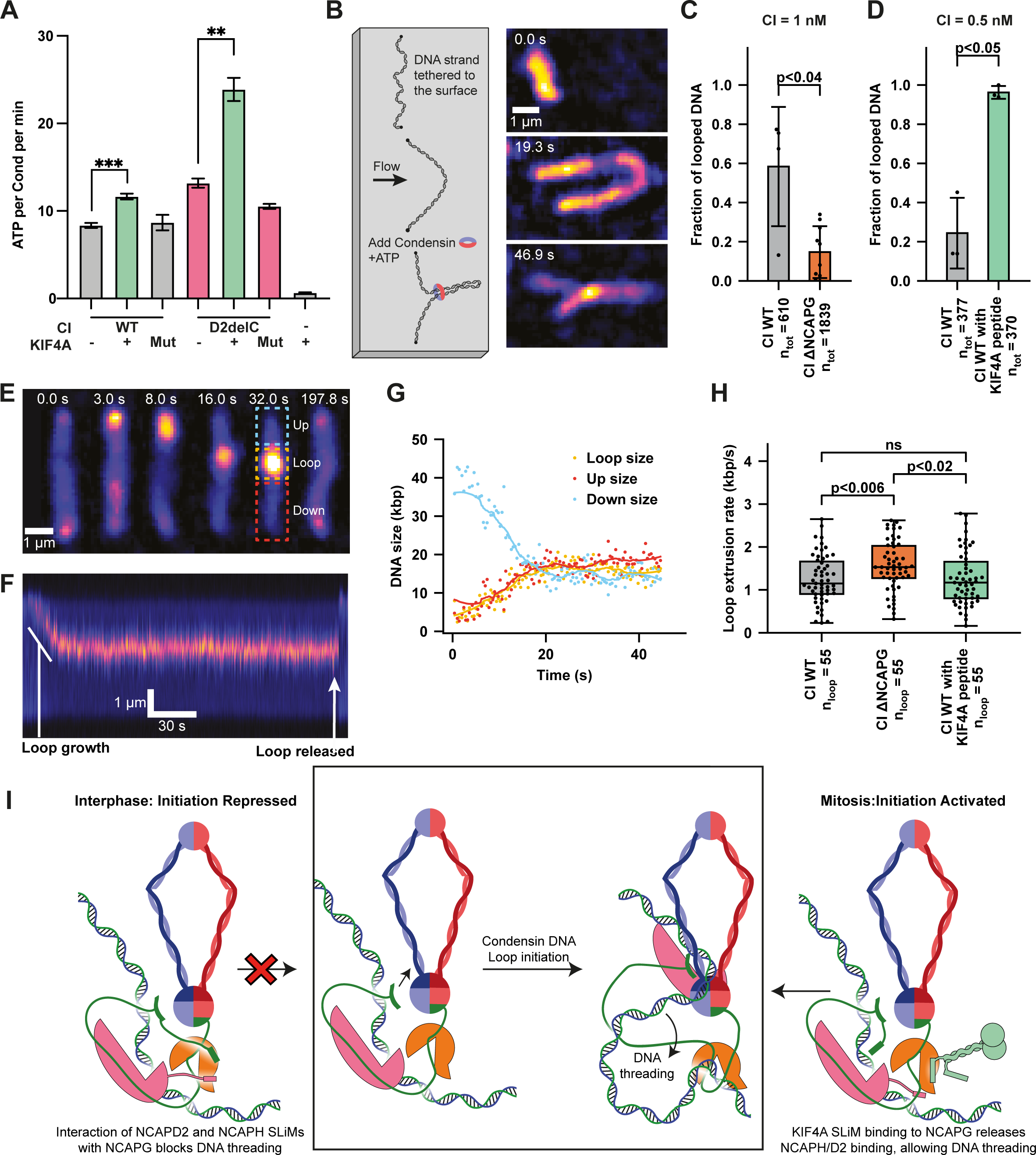
KIF4A increases condensin I loop initiation. (A) ATPase assay of WT and pentameric condensin I lacking the C-terminal region of NCAPD2 in the presence and absence of WT or mutant KIF4A_1206-1228_ peptide. ATPase assays from at least 3 repeats, error bars indicate standard error, analysed with two tail t-test. (B) Schematic of the single-molecule loop extrusion assay under constant buffer flow and corresponding snapshots of a loop formed by human Condensin I WT in the presence of KIF4A_1206-1228_ peptide. (C) and (D) Fraction of DNA strands (mean ± sd) with at least one looping event during a 1000 s acquisition time in the presence of 1 nM human Condensin I WT and ΔNCAPG, and 0.5 nM Condensin I WT and WT supplied with 1000x KIF4A_1206-1228_ peptide, respectively. (E) Example snapshots, the corresponding (F) kymograph and (G) time trace of DNA lengths for a loop extrusion event by the human Condensin I ΔNCAPG without flow. (H) Box-whisker-plots showing the loop extrusion rates of human Condensin I WT, ΔNCAPG and WT in the presence of KIF4A_1206-1228_ peptide. For the box plots, the central line denotes the medians, the box limit denotes the 25th–75th percentile and the whiskers denote 5th–95th percentile. The p values were calculated using Welch’s t-test. *p<0.05, **p<0.01, ***p<0.001, **** p<0.0001. (I) Model of KIF4A/condensin I activation mechanism. Central box illustrates loop initiation step, where DNA is threaded through the NCAPH Kleisin. In the presence of interactions between NCAPD2/H and NCAPG, the threading step is blocked (left), while interaction with KIF4A allows DNA threading and loop initiation (right).

### KIF4A promotes Condensin I loop initiation

As KIF4A was able to activate condensin I ATPase activity, and has previously been shown to activate condensin I in cells^25^, we next aimed to determine the effect of this interaction on condensin I loop extrusion. We used a well-established single-molecule assay for directly visualising loop extrusion by SMC complexes, whereby a 48.5 kbp piece of lambda DNA is flow stretched, attached to a passivated surface via streptavidin-biotin binding and stained with Sytox Orange (SxO) for fluorescence imaging^2^. DNA is visualised with total internal reflection microscopy and can be stretched perpendicular to its attachment axis by buffer flow (Figure 3B). After addition of 1 nM of condensin I and 2.5 mM ATP under flow a DNA punctum forms, which gradually increases in size over time confirming the event as DNA loop extrusion (Figure 3B, Supplementary Movie 1).

First, we tested tetrameric condensin I lacking NCAPG, and found it was able to create loops. However the proportion of tethers looped was less than for wild-type (Figure 3C), likely due to reduced DNA binding affinity (Figure 1D). Next, we repeated the assay in the presence or absence of 0.5 μM of KIF4A_1206-1228_ peptide, reducing the condensin I concentration to 0.5 nM. While wild-type condensin I looped ∼24% of tethers, addition of the peptide resulted in ∼96% of tethers being looped, suggesting KIF4A promoted loop formation (Figure 3D).

We quantified loop extrusion rate in the absence of flow (Supplementary Movie 2) by measuring how the length of DNA above and below the loop (up and down) changed over time (Figure 3E-G). While loss of NCAPG increased loop extrusion rate compared to the wild-type, addition of KIF4A_1206-1228_ peptide resulted in no significant difference (Figure 3H), consistent with the effects on ATP hydrolysis rate. This suggests KIF4A could promote loop initiation rather than affecting loop extrusion rate. Current mechanisms for loop extrusion suggest that loop initiation requires DNA to be “threaded” through a compartment formed by the kleisin. Interactions between the NCAPH/D2 with NCAPG could impede this threading process, inhibiting loop initiation. Binding of KIF4A would remove this blockage, allowing DNA threading and condensin I loop initiation (Figure 3I).

### Motif binding is conserved across species

We have shown that NCAPG acts as a scaffold, binding multiple SLiMs. The equivalent subunits in condensin II and cohesin, NCAPG2 and STAG1/2, respectively, also act as a scaffold for SLiM interactions^22, 34, 35^, hence we speculate that this is a conserved mechanism. Previous work on *C. thermophilum* condensin observed tetrameric condensin lacking NCAPG equivalent Ycg1 resulted in a higher ATPase rate^15^. We examined sequence data and found *C. thermophilum* NCAPH equivalent, Brn1, contained the N-terminal motif (Supplementary Figure 3A), suggesting this interaction is conserved outside metazoans.

While the C-terminal region of NCAPD2 is absent in yeast the residues it contacts on NCAPG are conserved (Supplementary Figure 3B and C), suggesting it may act as binding site for other factors. We searched literature for factors that have been reported to interact with yeast condensin and performed AlphaFold2 predictions. We found two proteins, Lrs4 and Sgo1, contained similar SLiMs and were predicted to interact with Ycg1 in the equivalent site to NCAPD2/KIF4A with NCAPG (Figure 4A, B and C, Supplementary Figure 3D and E). Fluorescence polarisation binding assays, indicate peptides of Lrs4_317-337_ and Sgo1_503-523_ bind with Kd of 12.9 ± 2.5 μM and 4.0 ± 0.3 μM to pentameric yeast condensin, respectively, while negligible binding to condensin lacking Ycg1 was observed (Figure 4D and E). Sgo1 condensin interaction mapping using peptide array support this region as being an interaction site, in addition to the Serine-Rich Motif spanning residues 137-163^36^, while mutagenesis of Lrs4 demonstrated that a Q325stop mutation resulted in synthetic lethality on a smc2-157 mutated background and exhibited shortening of rDNA^37^, providing additional support for the significance of these interactions and a general mechanism of regulation by SLiMs binding HAWKs (Figure 4F).

**Figure 4:**
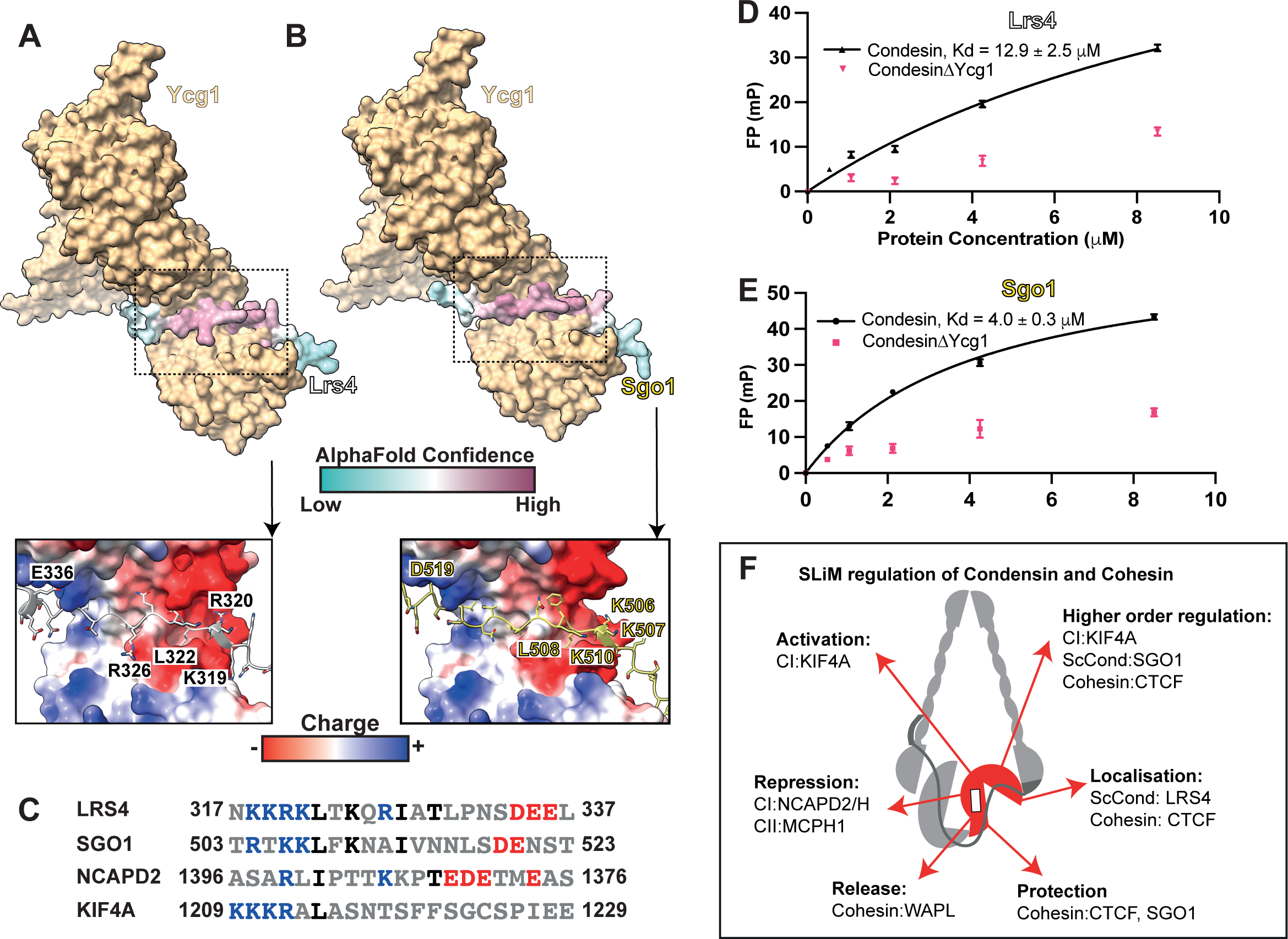
SLiM HAWK association is a general mechanism in condensin and cohesin. (A) and (B) AlphaFold2 model of Ycg1 with Lrs4 and Sgo, respectively, with insert of electrostatic potential. (C) Sequence alignment of SLiMs that bind the equivalent site of NCAPG and Ycg1. (D and E) Fluorescence polarisation binding data of Lrs4 and Sgo peptides, respectively, with yeast condensin complex. (F) A general model of the role SLiM HAWK association has in regulating condensin and cohesin activity.

## Discussion

KIF4A has long been implicated in chromosome formation with cellular data demonstrating that loss of condensin I/KIF4A association results in loss of condensin I loading, poorly condensed chromosomes and congregation defects^24, 25^. This work provides a mechanism to explain this phenotype, where KIF4A activates the condensin I complex by competing off auto-inhibitory interactions (Figure 3I).

Condensin I activation has previously been attributed to phosphorylation events, and while this work suggests they are not solely responsible, likely they contribute to it. The SLiMs of NCAPH, NCAPD2 and KIF4A have numerous phosphorylation sites (Supplementary Figure 4A). Three families of kinases are likely to play a role. Firstly, phosphorylation sites have been identified in NCAPD2 1366-1386, which are enriched in G1^38^. The sequence in this region suggests acidic directed kinases, such casein kinases, which have previously been suggested to phosphorylate NCAPD2 to inhibit condensin I during interphase^39^. Secondly, CDK1 activity peaks during mitosis and all three SLiMs identified contain conical S/TP CDK1 consensus motifs^40^. Finally, Aurora kinases have been implicated in KIF4A mediated condensin I activity. Inhibition of Aurora B kinase and mutation of Aurora consensus site in the NCAPH SLiM, S70A, reduces KIF4A and condensin I chromosome association^41^, while Aurora A activity is essential for KIF4A dependent chromosome congregation^25^. Aurora kinases have been demonstrated to phosphorylate both NCAPH S70 and NCAPD2 S1395^42, 43^ (Supplementary Figure 4A). Based on the AlphaFold2 models the surface charge suggests phosphorylation by casein kinase likely enhances NCAPD2/NCAPG association, aiding condensin I repression in G1, while CDK1 and Aurora phosphorylation results in charge clashes for inhibitory NCAPD2/NCAPH interactions with NCAPG, while adding complementary charge for the KIF4A interaction, likely aiding binding of KIF4A in mitosis. Adding to the complexity of regulation by phosphorylation is the role of phosphatases, as protein phosphatase 2 and 4 bind to KIF4A and NCAPD2, respectively^26, 44^. Future studies should address the effect and timing of addition and removal of these phosphorylations, and the spatial and temporal effect they have on condensin activity.

Our findings are highly complementary to previous studies, which demonstrated that loss of NCAPG still enabled chromosome formation while deletion or phosphomimic mutations of the N-terminus of NCAPH increased loop extrusion and chromosome condensation^20, 29^. Our loop extrusion rate of 1.2-1.5 kbp/s is comparable to that previously determined for human condensin I^3, 29^, however we see a greater enhancement of the fraction of tethers with loops on the addition of the KIF4A peptide than that observed on deletion of NCAPH N-terminus^29^, potentially illustrating that KIF4A’s actions involve competition with both the NCAPH and NCAPD2 SLiMs. Our data indicates the binding affinity of KIF4A to condensin I is relatively weak. Previous studies demonstrated the cysteine rich region of KIF4A is essential for condensin I association in cells^25^. This suggests an additional KIF4A/condensin I interaction site, which could result in binding avidity, hence higher affinity. However, fluorescence recovery experiments indicate KIF4A is more dynamic on chromosomes than condensin I^24^, suggesting KIF4A and condensin form a transient complex in cells.

The secondary finding of this work is the emerging pattern of the SMC complexes condensin and cohesin being finely regulated by the binding of SLiMs to the NCAPG equivalent HAWK subunits (Figure 4F). In this work, we have demonstrated that SLiMs from condensin I subunits NCAPD2 and H repress complex activity, while KIF4A activates it. SLiM mediated regulation at this site is conserved across species, with the N-terminal NCAPH SLiM being present in *C. thermophillium* Brn1 condensin subunit. This explains the previously observed result that *C. thermophillium* condensin lacking NCAPG equivalent HAWK, Ycg1, exhibits both higher ATPase activity and a higher proportion of DNA tethers being compacted in loop extrusion assays^15^. We have identified SLiMs which bind Ycg1 in the known yeast condensin binding partners Sgo and Lrs4. Future work should address the function of these interactions. The simplest hypothesis would be that they help facilitate recruitment, which is likely the case for Lrs4, where deletion of the SLiM results in shortening of rDNA^37^. However, binding could also alter temporal or spatial regulation, which may be the case for Sgo1, as the Ycg1 binding region (residues ∼500-520), is adjacent to APC/C D-box degron^45^. Previous studies have also demonstrated SLiM interactions regulate condensin II and cohesin. During interphase, condensin II is repressed by MCPH1 which binds via a SLiM to the NCAPG equivalent, NCAPG2^22^, while the cohesin NCAPG equivalent subunit, STAG1/2, is bound by SLiMs found in WAPL, promoting cohesin unloading, CTCF, resulting in stalling/localisation and SGO1, protecting centromeric cohesin^35, 46^. In many of these examples the binding of the SLiM is just one of multiple interactions, for example the cysteine rich region of KIF4A also necessary for condensin I binding in cells^25^, and avidity contributing to bind is a common feature of SLiM interactions^47^. Hence, this work and the work of others demonstrates that SLiMs binding HAWK subunits of the SMC complexes condensin and cohesin is a conserved mechanism, resulting in diverse regulatory outcomes (Figure 4F).

## Limitations of the study

This study uses AlphaFold2 multimer^32^ to understand the molecular mechanism underlying the KIF4A mediated activation of condensin I. While these predicted interactions are supported by activity and binding assays, as well as existing cellular studies, they have weak binding affinities and could enhanced by avidity generated from other interaction sites as well as post-translational modifications. However, weak binding affinity may be necessary to ensure dynamic and reversible changes in activity of these complex. Experimental, high-resolution structures would be needed to fully understand the details of interactions, both within the condensin I complex, and with cofactors, such as KIF4A.

## Acknowledgement

We thank David Rueda, Paul Girvan and Anita Meier for assistance in collecting proof of principal single molecule data, and Alessandro Vannini for human condensin I constructs.

This study was funded by the Medical Research Council UKRI MC-A652-5PY00 (L.A., and E.E.C), the Max Planck Society (EK), European Research Council Starting Grant 101076914 (EK), Deutsche Forschungsgemeinschaft (DFG, German Research Foundation) – SFB 1551 (DT, EK) and IMPRS on Cellular Biophysics (DT).

## Author Contribution

E. C. conceptualization, data curation, formal analysis, investigation, methodology, visualisation, writing-original draft. D.T. data curation, formal analysis, investigation, visualisation and writing-reviewing and editing. L. A. and E. K. Funding acquisition, resources, supervision, writing-reviewing and editing.

## References

1. Sundin, O., and Varshavsky, A. (1981). Arrest of segregation leads to accumulation of highly intertwined catenated dimers: Dissection of the final stages of SV40 DNA replication. Cell 25, 659–669. 10.1016/0092-8674(81)90173-2.

2. Ganji, M., Shaltiel, I.A., Bisht, S., Kim, E., Kalichava, A., Haering, C.H., and Dekker, C. (2018). Real-time imaging of DNA loop extrusion by condensin. Science 360, 102–105. 10.1126/science.aar7831.

3. Kong, M., Cutts, E.E., Pan, D., Beuron, F., Kaliyappan, T., Xue, C., Morris, E.P., Musacchio, A., Vannini, A., and Greene, E.C. (2020). Human Condensin I and II Drive Extensive ATP-Dependent Compaction of Nucleosome-Bound DNA. Mol. Cell, 683540. 10.1016/j.molcel.2020.04.026.

4. Gibcus, J.H., Samejima, K., Goloborodko, A., Samejima, I., Naumova, N., Nuebler, J., Kanemaki, M.T., Xie, L., Paulson, J.R., Earnshaw, W.C., et al. (2018). A pathway for mitotic chromosome formation. Science 359, eaao6135. 10.1126/science.aao6135.

5. Sonoda, E., Matsusaka, T., Morrison, C., Vagnarelli, P., Hoshi, O., Ushiki, T., Nojima, K., Fukagawa, T., Waizenegger, I.C., Peters, J.-M., et al. (2001). Scc1/Rad21/Mcd1 Is Required for Sister Chromatid Cohesion and Kinetochore Function in Vertebrate Cells. Dev. Cell 1, 759–770. 10.1016/S1534-5807(01)00088-0.

6. Hauf, S., Waizenegger, I.C., and Peters, J.-M. (2001). Cohesin Cleavage by Separase Required for Anaphase and Cytokinesis in Human Cells. Science 293, 1320–1323. 10.1126/science.1061376.

7. Davidson, I.F., Bauer, B., Goetz, D., Tang, W., Wutz, G., and Peters, J.M. (2019). DNA loop extrusion by human cohesin. Science 366, 1338–1345. 10.1126/science.aaz3418.

8. Kim, Y., Shi, Z., Zhang, H., Finkelstein, I.J., and Yu, H. (2019). Human cohesin compacts DNA by loop extrusion. Science 366, 1345–1349. 10.1126/science.aaz4475.

9. Rao, S.S.P., Huang, S.-C., Glenn St Hilaire, B., Engreitz, J.M., Perez, E.M., Kieffer-Kwon, K.-R., Sanborn, A.L., Johnstone, S.E., Bascom, G.D., Bochkov, I.D., et al. (2017). Cohesin Loss Eliminates All Loop Domains. Cell 171, 305–320.e24. 10.1016/j.cell.2017.09.026.

10. Higashi, T.L., Pobegalov, G., Tang, M., Molodtsov, M.I., and Uhlmann, F. (2021). A brownian ratchet model for dna loop extrusion by the cohesin complex. Elife 10, 1–35. 10.7554/eLife.67530.

11. Pradhan, B., Kanno, T., Umeda Igarashi, M., Loke, M.S., Baaske, M.D., Wong, J.S.K., Jeppsson, K., Björkegren, C., and Kim, E. (2023). The Smc5/6 complex is a DNA loop-extruding motor. Nature 616, 843–848. 10.1038/s41586-023-05963-3.

12. Onn, I., Aono, N., Hirano, M., and Hirano, T. (2007). Reconstitution and subunit geometry of human condensin complexes. EMBO J. 26, 1024–1034. 10.1038/sj.emboj.7601562.

13. Collier, J.E., Lee, B.-G., Roig, M.B., Yatskevich, S., Petela, N.J., Metson, J., Voulgaris, M., Gonzalez Llamazares, A., Löwe, J., and Nasmyth, K.A. (2020). Transport of DNA within cohesin involves clamping on top of engaged heads by Scc2 and entrapment within the ring by Scc3. Elife 9, 264–277. 10.7554/eLife.59560.

14. Shi, Z., Gao, H., Bai, X., and Yu, H. (2020). Cryo-EM structure of the human cohesin-NIPBL-DNA complex. Science 368, 1454–1459. 10.1126/science.abb0981.

15. Shaltiel, I.A., Datta, S., Lecomte, L., Hassler, M., Kschonsak, M., Bravo, S., Stober, C., Ormanns, J., Eustermann, S., and Haering, C.H. (2022). A hold- and-feed mechanism drives directional DNA loop extrusion by condensin. Science 376, 1087–1094. 10.1126/science.abm4012.

16. Kschonsak, M., Merkel, F., Bisht, S., Metz, J., Rybin, V., Hassler, M., and Haering, C.H. (2017). Structural Basis for a Safety-Belt Mechanism That Anchors Condensin to Chromosomes. Cell 171, 588–600.e24. 10.1016/j.cell.2017.09.008.

17. Hara, K., Kinoshita, K., Migita, T., Murakami, K., Shimizu, K., Takeuchi, K., Hirano, T., and Hashimoto, H. (2019). Structural basis of HEAT-kleisin interactions in the human condensin I subcomplex. EMBO Rep. 20, e47183. 10.15252/embr.201847183.

18. Lee, B.G., Rhodes, J., and Löwe, J. (2022). Clamping of DNA shuts the condensin neck gate. Proc. Natl. Acad. Sci. U. S. A. 119, 1–33. 10.1073/pnas.2120006119.

19. Bazile, F., St-Pierre, J., and D’Amours, D. (2010). Three-step model for condensin activation during mitotic chromosome condensation. Cell Cycle 9, 3243–3255. 10.4161/cc.9.16.12620.

20. Kinoshita, K., Kobayashi, T.J., and Hirano, T. (2015). Balancing acts of two HEAT subunits of condensin I support dynamic assembly of chromosome axes. Dev. Cell 33, 94–107. 10.1016/j.devcel.2015.01.034.

21. Yoshida, M.M., Kinoshita, K., Aizawa, Y., Tane, S., Yamashita, D., Shintomi, K., and Hirano, T. (2022). Molecular dissection of condensin II-mediated chromosome assembly using in vitro assays. Elife 11, 2003–2005. 10.7554/eLife.78984.

22. Houlard, M., Cutts, E.E., Shamim, M.S., Godwin, J., Weisz, D., Presser Aiden, A., Lieberman Aiden, E., Schermelleh, L., Vannini, A., and Nasmyth, K. (2021). MCPH1 inhibits Condensin II during interphase by regulating its SMC2-Kleisin interface. Elife 10, 1–41. 10.7554/eLife.73348.

23. Wood, J.L., Liang, Y., Li, K., and Chen, J. (2008). Microcephalin/MCPH1 associates with the condensin II complex to function in homologous recombination repair. J. Biol. Chem. 283, 29586–29592. 10.1074/jbc.M804080200.

24. Samejima, K., Samejima, I., Vagnarelli, P., Ogawa, H., Vargiu, G., Kelly, D.A., de Lima Alves, F., Kerr, A., Green, L.C., Hudson, D.F., et al. (2012). Mitotic chromosomes are compacted laterally by KIF4 and condensin and axially by topoisomerase IIα. J. Cell Biol. 199, 755–770. 10.1083/jcb.201202155.

25. Poser, E., Caous, R., Gruneberg, U., and Barr, F.A. (2019). Aurora A promotes chromosome congression by activating the condensin-dependent pool of KIF4A. J. Cell Biol. 219, 650937. 10.1083/jcb.201905194.

26. Wang, X., Garvanska, D.H., Nasa, I., Ueki, Y., Zhang, G., Kettenbach, A.N., Peti, W., Nilsson, J., and Page, R. (2020). A dynamic charge-charge interaction modulates PP2A:B56 substrate recruitment. Elife 9, 1–19. 10.7554/eLife.55966.

27. Takahashi, M., Wakai, T., and Hirota, T. (2016). Condensin I-mediated mitotic chromosome assembly requires association with chromokinesin KIF4A. Genes Dev. 30, 1931–1936. 10.1101/gad.282855.116.

28. Lee, B., Merkel, F., Allegretti, M., Hassler, M., Cawood, C., Lecomte, L., O’Reilly, F.J., Sinn, L.R., Gutierrez-Escribano, P., Kschonsak, M., et al. (2020). Cryo-EM structures of holo condensin reveal a subunit flip-flop mechanism. Nat. Struct. Mol. Biol. 27, 743–751. 10.1038/s41594-020-0457-x.

29. Tane, S., Shintomi, K., Kinoshita, K., Tsubota, Y., Yoshida, M.M., Nishiyama, T., and Hirano, T. (2022). Cell cycle-specific loading of condensin I is regulated by the N-terminal tail of its kleisin subunit. Elife 11, 1–23. 10.7554/ELIFE.84694.

30. Ball, A.R., Schmiesing, J.A., Zhou, C., Gregson, H.C., Okada, Y., Doi, T., and Yokomori, K. (2002). Identification of a Chromosome-Targeting Domain in the Human Condensin Subunit CNAP1/hCAP-D2/Eg7. Mol. Cell. Biol. 22, 5769– 5781. 10.1128/mcb.22.16.5769-5781.2002.

31. Jumper, J., Evans, R., Pritzel, A., Green, T., Figurnov, M., Ronneberger, O., Tunyasuvunakool, K., Bates, R., Žídek, A., Potapenko, A., et al. (2021). Highly accurate protein structure prediction with AlphaFold. Nature 596, 583–589. 10.1038/s41586-021-03819-2.

32. Evans, R., O’Neill, M., Pritzel, A., Antropova, N., Senior, A., Green, T., Žídek, A., Bates, R., Blackwell, S., Yim, J., et al. (2022). Protein complex prediction with AlphaFold-Multimer. bioRxiv 804, 2021.10.04.463034. 10.1101/2021.10.04.463034.

33. Takahashi, M., Wakai, T., and Hirota, T. (2016). Condensin I-mediated mitotic chromosome assembly requires association with chromokinesin KIF4A. Genes Dev. 30, 1931–1936. 10.1101/gad.282855.116.

34. Li, Y., Muir, K., Bowler, M.W., Metz, J., Haering, C.H., and Panne, D. (2018). Structural basis for Scc3-dependent cohesin recruitment to chromatin. Elife 7. 10.7554/eLife.38356.

35. García-Nieto, A., Patel, A., Li, Y., Oldenkamp, R., Feletto, L., Graham, J.J., Willems, L., Muir, K.W., Panne, D., and Rowland, B.D. (2023). Structural basis of centromeric cohesion protection. Nat. Struct. Mol. Biol. 30, 853–859. 10.1038/s41594-023-00968-y.

36. Yahya, G., Wu, Y., Peplowska, K., Röhrl, J., Soh, Y.-M., Bürmann, F., Gruber, S., and Storchova, Z. (2020). Phospho-regulation of the Shugoshin - Condensin interaction at the centromere in budding yeast. PLOS Genet. 16, e1008569. 10.1371/journal.pgen.1008569.

37. Johzuka, K., and Horiuchi, T. (2009). The cis Element and Factors Required for Condensin Recruitment to Chromosomes. Mol. Cell 34, 26–35. 10.1016/j.molcel.2009.02.021.

38. Dephoure, N., Zhou, C., Villén, J., Beausoleil, S.A., Bakalarski, C.E., Elledge, S.J., and Gygi, S.P. (2008). A quantitative atlas of mitotic phosphorylation. Proc. Natl. Acad. Sci. U. S. A. 105, 10762–10767. 10.1073/pnas.0805139105.

39. Takemoto, A., Kimura, K., Yanagisawa, J., Yokoyama, S., and Hanaoka, F. (2006). Negative regulation of condensin I by CK2-mediated phosphorylation. EMBO J. 25, 5339–5348. 10.1038/sj.emboj.7601394.

40. Songyang, Z., Blechner, S., Hoagland, N., Hoekstra, M.F., Piwnica-Worms, H., and Cantley, L.C. (1994). Use of an oriented peptide library to determine the optimal substrates of protein kinases. Curr. Biol. 4, 973–982. 10.1016/S0960-9822(00)00221-9.

41. Poonperm, R., Takata, H., Uchiyama, S., and Fukui, K. (2017). Interdependency and phosphorylation of KIF4 and condensin I are essential for organization of chromosome scaffold. PLoS One 12, 1–15. 10.1371/journal.pone.0183298.

42. Lipp, J.J., Hirota, T., Poser, I., and Peters, J.-M. (2007). Aurora B controls the association of condensin I but not condensin II with mitotic chromosomes. J. Cell Sci. 120, 1245–1255. 10.1242/jcs.03425.

43. Sardon, T., Pache, R.A., Stein, A., Molina, H., Vernos, I., and Aloy, P. (2010). Uncovering new substrates for aurora a kinase. EMBO Rep. 11, 977–984. 10.1038/embor.2010.171.

44. Ueki, Y., Kruse, T., Weisser, M.B., Sundell, G.N., Larsen, M.S.Y., Mendez, B.L., Jenkins, N.P., Garvanska, D.H., Cressey, L., Zhang, G., et al. (2019). A Consensus Binding Motif for the PP4 Protein Phosphatase. Mol. Cell 76, 953–964.e6. 10.1016/j.molcel.2019.08.029.

45. Eshleman, H.D., and Morgan, D.O. (2014). Sgo1 recruits PP2A to chromosomes to ensure sister chromatid bi-orientation during mitosis. J. Cell Sci. 127, 4974–4983. 10.1242/jcs.161273.

46. Li, Y., Haarhuis, J.H.I., Sedeño Cacciatore, Á., Oldenkamp, R., van Ruiten, M.S., Willems, L., Teunissen, H., Muir, K.W., de Wit, E., Rowland, B.D., et al. (2020). The structural basis for cohesin–CTCF-anchored loops. Nature 578, 472–476. 10.1038/s41586-019-1910-z.

47. Holehouse, A.S., and Kragelund, B.B. (2023). The molecular basis for cellular function of intrinsically disordered protein regions. Nat. Rev. Mol. Cell Biol. 10.1038/s41580-023-00673-0.

48. Ashkenazy, H., Abadi, S., Martz, E., Chay, O., Mayrose, I., Pupko, T., and Ben-Tal, N. (2016). ConSurf 2016: an improved methodology to estimate and visualize evolutionary conservation in macromolecules. Nucleic Acids Res. 44, W344–W350. 10.1093/nar/gkw408.

49. Martínez-García, B., Dyson, S., Segura, J., Ayats, A., Cutts, E.E., Gutierrez-Escribano, P., Aragón, L., and Roca, J. (2022). Condensin pinches a short negatively supercoiled DNA loop during each round of ATP usage. EMBO J., 2022.06.03.494647. 10.15252/embj.2022111913.

50. Voulgaris, M., and Gligoris, T.G. (2019). A Protocol for Assaying the ATPase Activity of Recombinant Cohesin Holocomplexes. Methods Mol. Biol. 2004, 197–208. 10.1007/978-1-4939-9520-2_15.

