## Supplementary figures for "Molecular Mechanism of Condensin I Activation by KIF4A"

**Supplementary Figure 1: Condensin pentamers can be reconstituted from tetramers.** (A) and (B) Mass photometry data of condensin I and II tetramers lacking NCAPG or NCAPD2 respectively with and without sub-stoichiometric HAWK being added.

**Supplementary Figure 2: NCAPG peptide AlphaFold2 confidence.** (A), (B), (C) AlphaFold2 predicted alignment error, IDDT score per residue and SLiM region with conservation colouring from ConSurf[48], for NCAPG with NCAPD2, NCAPH and KIF4A peptide, respectively.

**Supplementary Figure 3: Condensin disordered sequence conservation.** (A) Alignment of N-terminal region of NCAPH/H2 and Brn1, in *H. sapiens, S. cerevisiae* and *C. thermophilium*, with NCAPG/Ycg1 and SMC2 binding regions indicated. (B) Alignment of C-terminal region of NCAPD2/3 and Ycs4 in *H. sapiens, S. cerevisiae* and *C. thermophilium.* (C) Alignment of SLiM docking region in NCAPG and Ycg1, from *H. sapiens and S. cerevisiae*, * indicates residues with in 4 Å of predicted bound SLiM. (D) and (E) AlphaFold2 predicted alignment error and IDDT score per residue for Ycg1 with Lrs4 and Sgo1, respectively.

**Supplementary Figure 4: Phosphorylation and phosphatase sites in SLiMs.** (A) Sites of phosphorylation by Aurora (orange triangle), CDK1/cyclin B (green circle) and casein (blue square) kinases in SLiM of NCAPD2, NCAPH and KIF4A. Red rectangles indicate phosphatase binding sites. (B), (C) and (D) The location of phosphorylation sites indicated in (A) in NCAPD2, NCAPH and KIF4A, respectively. Based off of electrostatic surface charge, the effect of the phosphorylation is coloured either green, for potentially enhancing the interaction or orange, for resulting in a potential charge clash.

**Supplementary Movie 1: DNA Loop extrusion by human Condensin I WT in the presence of KIF4A$_{1206-1228}$ peptide, with constant buffer flow.** Corresponding to Figure 3B.

**Supplementary Movie 2: DNA Loop extrusion by human Condensin I ΔNCAPG without buffer flow.** Corresponding to Figure 3E-H.

**Methods:**

| **Supplementary Table 1** | | |
|---|---|---|
| **Construct** | **Description** | **citation** |
| pBIG2abc Condensin I strep | | [3] |
| pBIG2abc Condensin I Q strep | SMC2 Q147L, SMC4 Q229L | [3] |
| pBIG2abc Condensin I delG strep | Lacking NCAPG | This work |
| pBIG2ab Condensin I delD2 strep | Lacking NCAPD2 | This work |
| pBIG2ab Condensin I Q delG ystrep | Lacking NCAPG with mutations: SMC2 Q147L, SMC4 Q229L | This work |
| pBIG2ab Condensin I Q delD2 ystrep | Lacking NCAPD2 with mutations: SMC2 Q147L, SMC4 Q229L | This work |
| pLIB NCAPG ystrep | | This work |
| pLIB NCAPD2 ystrep | | This work |
| pLIB NCAPG delC ystrep | Deletion after 913 | This work |
| pLIB NCAPD2 delC ystrep | Deletion after 1305 | This work |
| pLIB NCAPG patch1 ystrep | Mutations: D413K, E416K, E417K, R420D, K421D | This work |
| pLIB NCAPG patch 2 ystrep | Mutations: Q460A, E464K, S467A, E468K | This work |
| pLIB NCAPG acidic patch ($G_{ap}$) | D136K, D137K, D141K | This work |
| pLIB NCAPH delSB | Deletion of 421-539 | This work |
| pLIB NCAPH delN | Deletion 1-77 | This work |
| ScCondensin Pentamer | | [28,49] |

| | | |
|---|---|---|
| ScCondensin Tetramer | Lacking Ycg1 | [28,49] |
| **Peptides** | | |
| F-KIF4A WT$_{1206-1228}$ | 5FAM-PGKKKKRALASNTSFFSGCSPIE | This work |
| KIF4A WT$_{1206-1228}$ | PGKKKKRALASNTSFFSGCSPIE | This work |
| KIF4A Mut$_{1206-1228}$ | PGKKAAAAGAAATSFFSGCSPIE | This work |
| F-NCAPD2-C$_{1376-1398}$ | 5FAM-SAEMTEDETPKKTTPILRASARR | This work |
| F-N-NCAPH$_{55-77}$ | 5FAM-FPQNDDEKERLQRRRSRVFDLQF | This work |
| F-Lrs4$_{317-339}$ | 5FAM-NKKRKLTKQRIATLPNSDEEL | This work |
| F-Sgo1$_{503-523}$ | 5FAM-TRTKKLFKNAIVNNLSDENST | This work |
| **DNA oligos** | | |
| 50 Forward | GGTGTGACAGGGTGTGACAGGGTGTGACAGGG TGTGACAGGGTGTGACAG | [22] |
| Cy5 50 Reverse | /5Cy5/CTGTCACACCCTGTCACACCCTGTCACA CCCTGTCACACCCTGTCACACC | [22] |
| 50 Reverse | CTGTCACACCCTGTCACACCCTGTCACACCCT GTCACACCCTGTCACACC | [22] |

**Protein purification**

Constructs used for expression are in Supplementary Table 1. Mutations of NCAPH, NCAPD2 and NCAPG were generated using PCR, gel extracted and cloned using Gibson assembly (NEB, Gibson Assembly Master Mix, E2611). Incorporation of mutation was confirmed with Sanger Sequencing (Genewiz).

Human condensin pentamers and tetramers were purified as previously described in [3,22]. Briefly, wild-type and mutation of pentamer and tetramer and individual subunits NCAPG and NCAPD2 were cloned into biGBac vectors shown in Supplementary Table 1. Bacmids were generated by Tn7 transposition in DH10EMBacY cells (Geneva biotech), which were transfected into Sf9 cells with Cellfectin (GIBCO) to generate baculovirus. Virus was further amplified in Sf9s cells and each construct was expressed in ~500mL using either Sf9 or High Five insect cells, harvesting 72 hours after infection. Cells pellets were lysed in human condensin purification buffer (20 mM HEPES pH 8, 300 mM KCl, 5 mM MgCl2, 1 mM DTT, 10 % glycerol) supplemented with 1 Pierce protease inhibitor EDTA- free tablet (Thermo Scientific) or 1 cOmplete protease inhibitor EDTA-free tablet (Roche) per 50 mL and 25 U/ml of Benzonase (Sigma) with a Dounce homogenizer followed by brief sonication. Cleared lysate was loaded onto a StrepTrap HP or StrepTrap XT (Cytiva), washed with purification buffer before being eluted with purification buffer supplemented with 10 mM desthiobiotin or 50 mM Biotin (Sigma), respectively. Eluted fractions were diluted ~2 fold, before being loaded onto a Heparin HiTrap (cytiva) equilibrated with Heparin buffer A (20mM HEPES pH 8, 5% glycerol, 0.5mM DTT) with 150mM NaCl and eluted either via a gradient or step elution using Heparin buffer B (Heparin buffer A supplemented with 1M NaCl). Size exclusion chromatography was performed using human purification buffer on a Superose 6 16/60 (cytiva) for pentamers or tetramers or Superdex 200 30/10 column (cytiva) for individual HAWK protein. Protein containing fractions separated from the void volume were pooled, concentrated and flash frozen.

Yeast condensin pentameric and tetrameric complex were purified as described previously[28,49].

**Mass Photometry**

Mass photometry was used to demonstrate pentamer reconstitution from tetramer using a Refeyn TwoMP. A Refeyn sample carrier slide (Refeyn, MP-CON-21009) was mounting on the sample stage with a fresh silicon CultureWellTM gaskets (GBL103250, Sigma-Aldrich) attached to centre. All samples were measured in 20 mM HEPES pH 8, 150 mM KCl buffer using a field of view 512 x 138 pixels, collecting 6000 frames with a collection time of 60s. The focal position and imaging conditions were set using a 14 µL buffer droplet and data was collected by adding 2 µL of condensin sample, resulting in a final protein concentration of ~7 nM. All data were acquired with using the Refeyn AcquireMP software and analysed using the Refeyn DiscoverMP software. Masses were calibrated using the NativeMarkTM unstained protein standard (LC0725, Thermo Scientific) to generate a calibration curve

**ATPase activity assays**

ATPase assays were performed based on the method described previously[50] using the EnzCheck phosphate assay kit (Invitrogen). Condensin I complexes were used at a final concentration of 50 nM, while HAWK subunits were used at 100 nM and peptides were used at 50 µM. Basal ATPase assays were performed with final salt concentration of 100 mM KCl. DNA titrations were performed with 50bp dsDNA annealed in annealing buffer (10 mM Tris pH 7.5, 50mM NaCl, 1mM EDTA).

**EMSA**

The DNA sequence used for ATPase assay was used for gel shift assays, except a Cy5 labelled was added to the 5' of the reverse oligonucleotide (Supplementary Table 1). DNA was used at a final concentration of 50 nM, with indicated concentration of protein in buffer 20 mM HEPES pH 8, 150 mM KCl, 2.5 mM $MgCl_2$, 10% glycerol, 1 mM DTT. Sample was incubated on ice for 20 minutes before being resolved on a 2% agarose gel in 0.5x Tris borate (TB) buffer and scanned on a FUJIFILM FLA-5100 scanner.

**AlphaFold2 Predictions**

AlphaFold2 multimer predictions were performed using AlphaFold v2.3.1 using default parameters of the following Colab notebook with a Colab Pro account: https://colab.research.google.com/github/deepmind/alphafold/blob/main/notebooks/AlphaFold.ipynb

https://colab.research.google.com/github/sokrypton/ColabFold/blob/main/AlphaFold2.ipynb

Predictions of human condensin complex used residues 1-912 of NCAPG with 1-101 of NCAPH, 1307-1401 of NCAPD2 or 1065-1232 of KIF4A. Prediction of yeast condensin interaction used residue 1-475 of Ycg1 with full length Sgo1 or Lrs4.

Plots showing per residue pLDDT and per residue aligned error were generated using output PDB and JSON file using Python run in Jupyter notebooks. Figures of structural models were generated with ChimeraX, with regions of low prediction confidence hidden for clarity.

**Fluorescence Polarisation Assay**

Fluorescence polarisation binding assays were performed by mixing 100 nM of 5-FAM labelled peptide (Supplementary Table 1) with indicated concentration of protein in FP buffer (20 mM HEPES pH8, 150 mM KCl, 2.5 mM MgCl, 5% glycerol, 1 mM DTT) and incubated at room temperature for 15 minutes before reading the plate with a POLARstar Omega plate reader (BMG). The plate was read three times at 5 minute intervals to ensure samples reached equilibrium and three replicates were performed. Data was analysed in Graphpad Prism and Kd fit using the following:

$$FP = \left(\frac{FP_{max}}{2[Pep]}\right)\left(([C] + [Pep] + K_d) - \sqrt{([C] + [Pep] + K_d)^2 - 4.[C].[Pep]}\right)$$

Where [C] and [Pep] are the concentration, in μM, of condensin and 5-FAM labelled peptide respectively, $FP_{max}$ the maximum change in fluorescence polarization in mP and $K_d$ is the equilibrium dissociation constant. Fit curves are plotted, with standard error.

**Single molecule assay DNA substrate preparation**

The 48.5 kbp DNA substrate used in the single molecule assay in this study was prepared as follows. Linear and double-stranded (48502 bp) Lambda Phage DNA with 12 bp single-stranded 5'-ends was purchased from New England Biolabs (N3011L). The complementary biotinylated primers were ligated with Taq DNA Ligase to achieve biotinylation on both ends of the Lambda DNA. The Lambda DNA biotinylated at both ends was purified from the 10x molar excess primers and the ligase by a custom built Äkta column.

**Single molecule Flow cell Preparation**

The microscope slides (76x26x1 mm microscope slides Marienfeld 1000000) and coverslips (24x60 mm No1.5 thickness 170 μm, borosilicate VWR Intl 631-0147) used in this study were functionalized as previously described (Chandradoss et al., 2014) with the minor following modifications. Microscope slides were laser drilled to achieve 11 channels to attach inlet / outlet tubings and reused several times. Since an objective-type TIRF microscope was used the coverslips were also treated with 1 M of KOH solution and etched with acid Piranha solution of 5:1 sulfuric acid:hydrogen peroxide ratio prior to amino-silanization. The microscope slides and coverslips were silanized with a solution of 100 mL of anhydrous methanol, 5 mL of acetic acid and 10 mL of APTES ((3-aminopropyl)triethoxysilane) and PEGylated with 4 mg of m-PEG-SVA (MW 5000 Laysan Bio) and 0.1 mg Biotin-PEG-SVA (MW 5000 Laysan Bio) in 50 mM Boric Acid, 12.5 mM Sodium Hydroxide pH 8.5. The PEGylation was repeated 4 times for at least 4 hours at 4°C. The microscope slides and coverslips were washed with MQ water and dried with a gentle flow of nitrogen between the PEGylation steps.

After the 4th PEGylation the microscope slides and coverslips were sealed and stored at -20°C until the assembly of the flow cell. Prior to the flow cell assembly, a 5th PEGylation step using 50 mM MS(PEG)4 (Thermo Scientific) in 50 mM Boric Acid, 12.5 mM Sodium Hydroxide pH 8.5 was done overnight at 4°C. The shorter (MW 333) PEG molecules were used in the last PEGylation step since they may be more effective in filling up possible holes in the PEG layer. After this step the microscope slide and coverslip were washed with MQ water, dried with a gentle flow of nitrogen and used directly in the assembly of a flow cell.

Multi-channel flow cells were assembled using the functionalized microscope slides and coverslips with double sided scotch tape as a spacer between the two to achieve 2 mm x 24 mm x 100 μm channels. The channels were sealed with Epoxy glue. The outlets were assembled using 1-200 μl Axgen pipette tips, tubings and 0.5x10 mm syringe needles glued together with Epoxy, attached to the laser drilled holes of the microscope slide and glued in place with Epoxy. 1-200 μl Axygen pipette tips were used as inlet during experimenting.

##### HiLo microscopy and data acquisition

The HiLo microscope used in this study was described previously [11].

##### Single-molecule loop extrusion assay

Real time imaging of the loop extrusion by Condensin I was carried out in the aforementioned flow cells as follows. The channel was washed with T50 buffer (40 mM Tris-HCl pH8.0, 50 mM NaCl, 0.2mM EDTA), incubated with 1 μM Streptavidin in T50 buffer for 2 minutes and incubated with 5 mg/ml BSA in T50 buffer for 5 minutes to further reduce unspecific binding of proteins to the surface. The excess Streptavidin and BSA were washed thoroughly with T50 buffer. Lambda DNA biotinylated at both ends was introduced to the channel at a flow rate of 2 μl/min facilitating surface attachment through the Streptavidin-Biotin binding. The flow was adjusted and kept constant with a syringe pump attached to the outlet of the flow cell. Excess DNA was washed thoroughly with T50 buffer.

The surface attached DNA molecules were imaged in the imaging buffer (40 mM Tris-HCl pH 7.5, 2 mM Trolox, 30 mM D-glucose, 50 mM Potassium Glutamate (L-Glutamic acid monopotassium salt monohydrate), 2.5 mM MgCl2, 1 mg/ml BSA, 1 mM TCEP, 2.5 mM ATP, 100 nM Sytox Orange, 30 mg/ml Glucose Oxidase (15 U/ml), 20 mg/ml Catalase (1000 U/ml)). The imaging buffer supplied with used Condensin I variant was introduced into the channel at a flow rate of 10 µl/min for 100s and the flow was stopped afterwards. For the activity assay comparing Condensin I WT and Condensin I ΔNCAPG 1 nM of protein, for the activity assay comparing Condensin I WT and Condensin I WT supplied with KIF4A peptide 0.5 nM of protein and 1000x of the KIF4A peptide, for the analysis of loop extrusion rate 0.75 nM – 3 nM of protein (in the case of Condensin I WT supplied with KIF4A peptide, correspondingly 1000x of KIF4A peptide) was used.

**Single-molecule Data analysis**

A custom-written Python code was used to analyze the fluorescence image series. (Pradhan et al., 2023b) Python packages Napari (https://github.com/Napari/napari) and PyQtGraph (https://github.com/pyqtgraph/pyqtgraph) were used for the visualization and evaluation of the data.

**Analysis of loop extrusion activity**

Region of interests (ROIs) containing a single Lambda DNA molecule were chosen manually and annotated using the tools provided by the Python package Napari. For a single loop extrusion experiment a field of contains approximately 100-200 DNA strands. Only the strands that were tethered at both ends from the beginning to the end of the data acquisition were considered for the analysis. Each of the DNA strands were manually analyzed whether they form at least one loop during an acquisition time of 1000 s, including the initial 100 s of flow. The “fraction of looped DNA” (Figure B and C) was defined as the ratio of number of the DNA strands which form at least one loop to

the total number of DNA strands. This quantity was taken as a measure for the activity of the protein. For the average fraction of looped DNA (Figure B and C) the mean of multiple, independent experiments was taken. (For 1 nM Condensin I WT 4, for 1 nM

Condensin I ΔNCAPG 10, for 0.5 nM Condensin I WT 3, For 0.5 nM Condensin I WT supplied with 0.5 µM KIF4A 3 independent experiments were performed and analyzed.) The error bars in the figure are given by the standard deviation of these datasets. Since the number of independent data points for each of the variants was not sufficient to assume normality of the distribution we applied the non-parametric Wilcoxon rank-sum test to determine the p-values. The stats package from the Python module scipy was used for the statistical tests.

**Analysis of loop extrusion rate**

ROIs with a DNA strand that formed a loop were cropped and saved in TIFF format for further analysis. For snapshots shown in the figures (A and D) a median filter with a radius of 2 pixels was applied for smoothing the image. Furthermore, a white top-hat filter with radius 10 was applied for background subtraction using the "white_tophat" function from the package ndimage from the Python module scipy. The kymographs were built by summing the fluorescence intensity for 11 pixels along the line centered around the DNA axis from the median filtered images. Each vertical line in the kymograph corresponds to one frame of the image series.

For the determination of the loop extrusion rate, each line in the kymograph was split into three regions: "Up", "Loop" and "Down" (Figure D). For this, first the loop region was determined as follows: Using the "find_peaks" function from the package signal from the Python module scipy local intensity maxima in each line of the kymograph was determined. The center position of the DNA loop was chosen as the most intense peak within one line. The single peaks along the kymograph were connected using the "link" function from the Python package trackpy while manually supervising that the connected peaks correspond to the extruded loop. The region "Loop" was then assigned to the 9 pixels along the center peak for each line of the kymograph. The regions "Up" and "Down" were assigned to the remaining upper and lower portion of the kymograph, respectively for each line. The amount of DNA in each of the regions was estimated by multiplying the ratio of fluorescence intensity of each region to the total fluorescence intensity with 48.5 kbp which corresponds to the total length of the used Lambda DNA strands. The DNA size in each region against time gives the kinetics of a loop formation. In Figure 3G the kinetics of an exemplary loop is plotted together with the smoothed data using a Savitzky-Golay second order filter and 50 points window size. The loop extrusion rate in Fig. 3H was determined by fitting a linear function $f(x)=kx+c$ to the initial 5 seconds of the loop growth curve. The 5 second window of the fitting was chosen after the initiation of the loop where the loop size was increasing monotonously. Loop formation events were left out from the analysis of the loop extrusion rate if the signal was not clear enough which could be due to several reasons, eg. other DNA strands in the way, the loop formation event is not very stable and the loop slides back within the initial 5 second window, the loop forms at the upper or lower edge of the kymograph making it not possible to determine the size of the size of the “Up” or “Down” portion, respectively etc.

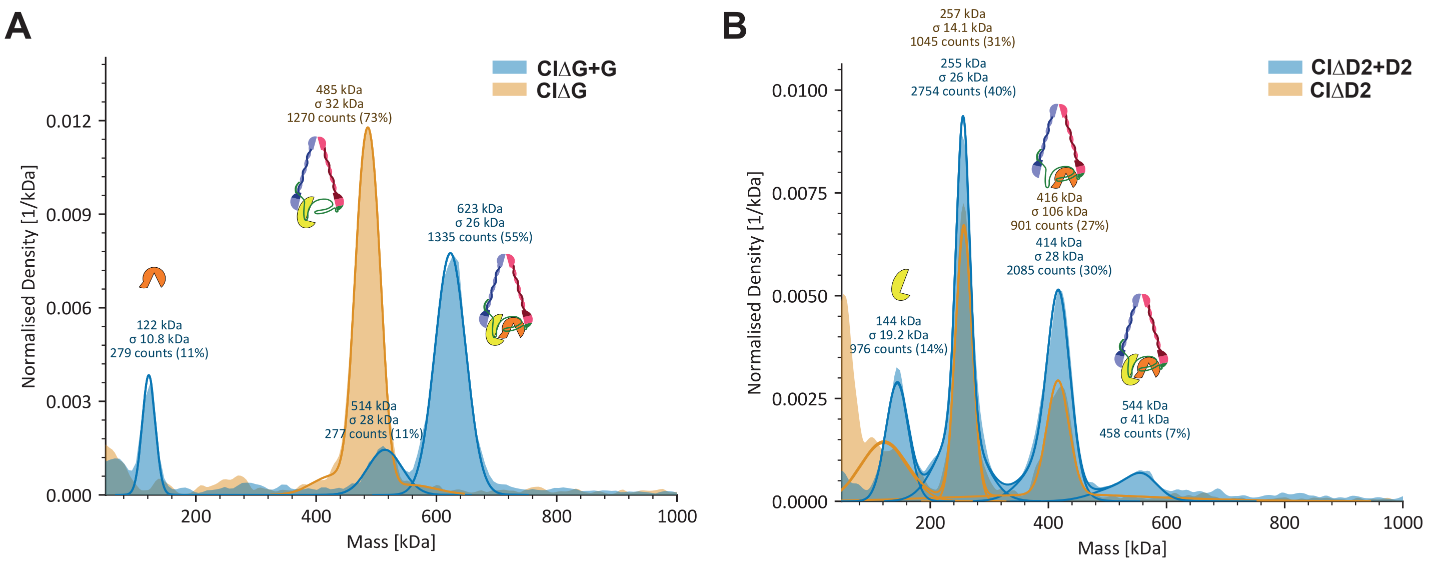


### Supplementary Figure 1

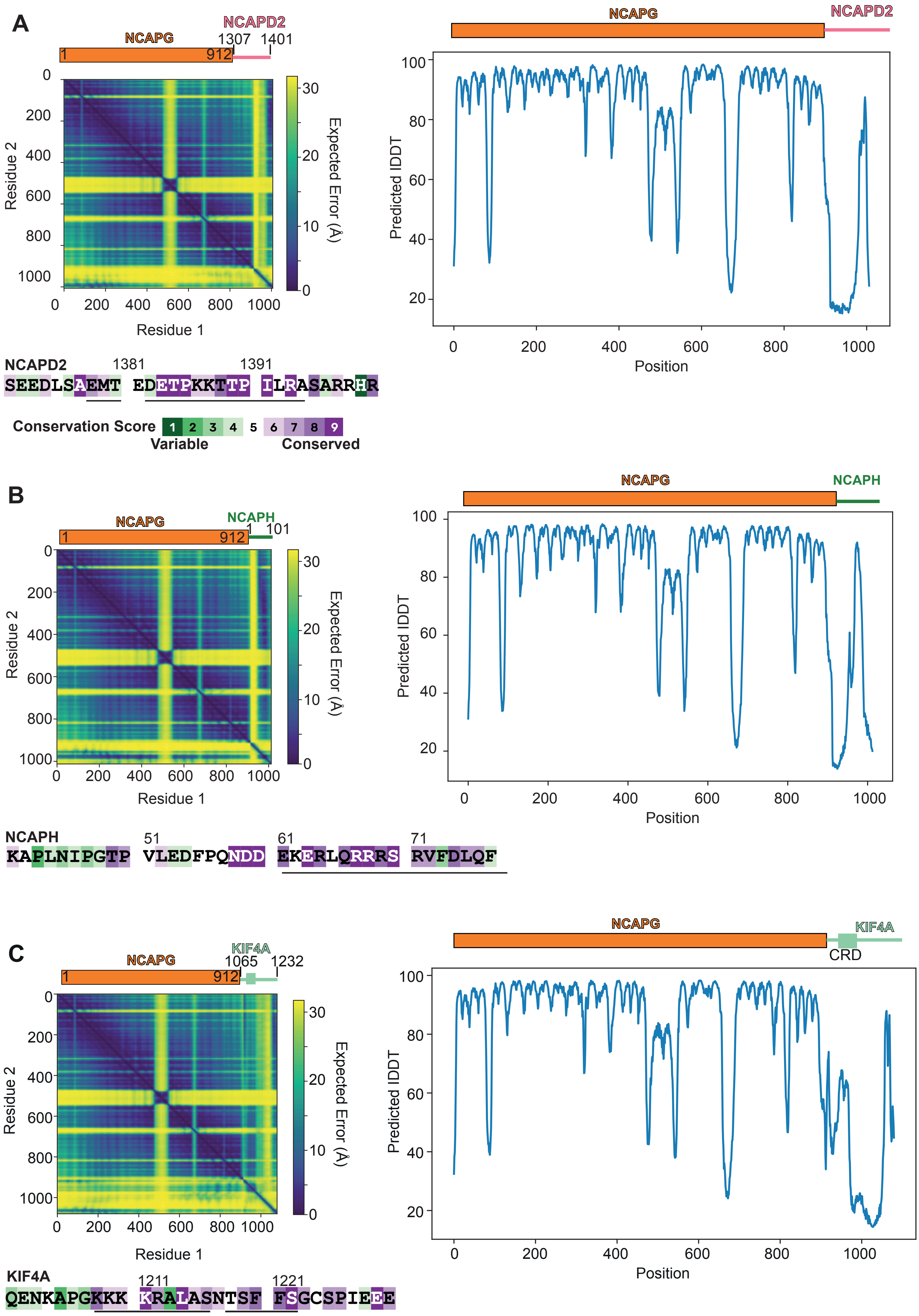


### Supplementary Figure 2

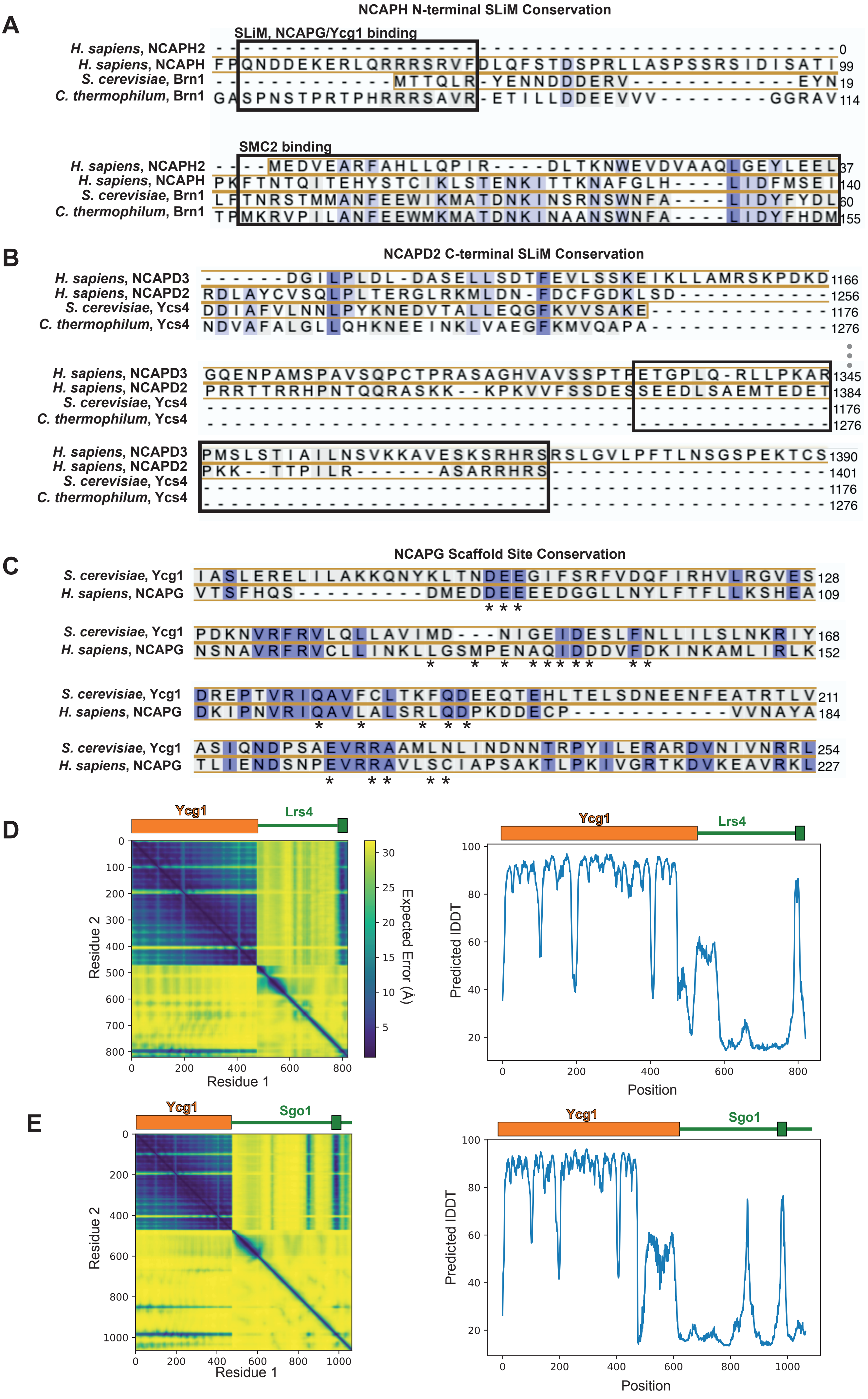


**Supplementary Figure 3**

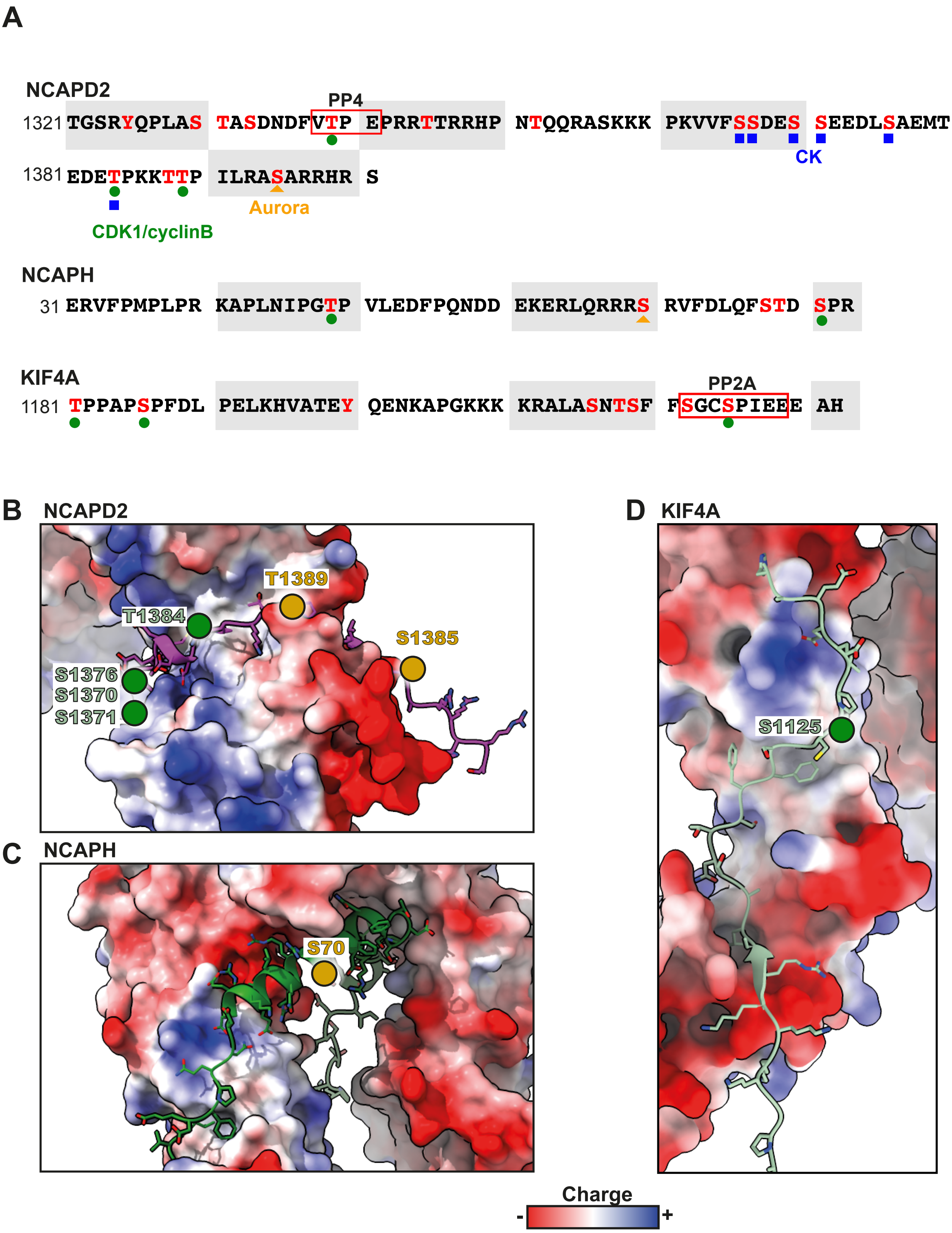


### Supplementary Figure 4
